# Genomic analyses identify significant molecules and biological processes in colorectal cancer cells with DNA damage

**DOI:** 10.1101/2022.01.24.477593

**Authors:** Hongmei Guo, Mason Zhang, Hanming Gu, James Liu

**Author notes:** Corresponding author: Hanming Gu, SHU-UTS SILC School, Shanghai University, Shanghai, China.

## Abstract

Colorectal cancer is a major cause of cancer deaths in the US. DNA damage is considered to be a novel target for the treatment of colorectal cancer. However, the molecular mechanisms and functions are still unclear. In this study, we aim to identify the significant molecules and signaling by analyzing the RNA-seq data. The GSE189366 was created by the BGISEQ-500 (Homo sapiens). The KEGG and GO analyses indicated the p53 signaling pathway and Hippo signaling pathway are major affected processes in colorectal cancer by DNA damage. Furthermore, we identified ten key interactive molecules including CDK1, STAT3, MDM2, CCNB1, CCNA2, CDKN1A, PCNA, AURKA, PLK1, and CDC6. Our study may provide potential drug targets for colorectal cancer.

## Introduction

Colorectal cancer is the third most cancer in the US^1^. The total incidence of colorectal cancer has decreased, but the incidence of colon and rectal cancer in young people has increased^2^. In recent studies, male sex and aging adult show a strong relationship with colorectal cancer^3^. In addition, hereditary and environmental risks are involved in the development of cancer progression^4^. Positive family history showed 10-20% of patients with colorectal cancer^5^. Young adult colorectal cancers are related to hereditary colorectal cancer syndromes and the reason is still not clear^6^. Though a number of genome studies have identified cancer susceptibility genes that are closely related to colorectal cancer risk, most factors are still unclear and subject to further study.

Colorectal cancer is caused by a complex of molecular events, for example, the mismatch repair alterations account for 5-15% of all colorectal cancer patients^7^. Colorectal cancer contains two subtypes: microsatellite unstable or microsatellite stable^8^. One study showed that patients with microsatellite stable cancers can be treated by the checkpoint inhibitors such as pembrolizumab or nivolumab^9^. With the need for better improvement, genomic alterations in the DNA damage response pathway are emerging as novel targets for treatment across different cancers^10^.

In our study, we analyzed the impacts of DNA damage on colorectal cancer by using the RNA-seq data. We identified a number of DEGs and functional biological processes. We also performed the gene enrichment and created the protein-protein interaction (PPI) network to figure out the relationships among the DEGs.

## Methods

### Data resources

Gene dataset GSE189366 was obtained from the GEO database. The data was produced by the BGISEQ-500 (Homo sapiens) (Cancer Biology Laboratory, Molecular Biology, Ariel University, RamatHa Golan Street 65’, Ariel, Israel). The analyzed dataset includes 2 of controls and 6 of topoisomerase (Top1) inhibitor irinotecan treated HCT116 cell lines.

### Data acquisition and processing

The data were organized and conducted by R package as previously described^11–14^. We used a classical t-test to identify DEGs with P< 0.001 and fold change ≥1.5 as being statistically significant.

The Kyoto Encyclopedia of Genes and Genomes (KEGG) and Gene Ontology (GO) KEGG and GO analyses were conducted by the R package (ClusterProfiler) and Reactome. P<0.05 was considered statistically significant.

### Protein-protein interaction (PPI) networks

The Molecular Complex Detection (MCODE) was used to create the PPI networks. The significant modules were produced from constructed PPI networks and String networks. The pathway analyses were performed by using Reactome (https://reactome.org/), and P<0.05 was considered significant.

## Results

### Identification of DEGs in colorectal cancer after DNA damage by irinotecan

To determine the effects of DNA damage and mutations on cancer cells, we analyzed the RNA-seq data from colorectal cancer with irinotecan treatment. A total of 1010 genes were identified with the threshold of P < 0.0001. The top increased and decreased genes were indicated by the heatmap and volcano plot (Figure 1). The top ten DEGs were listed in Table 1.

**Figure 1.**
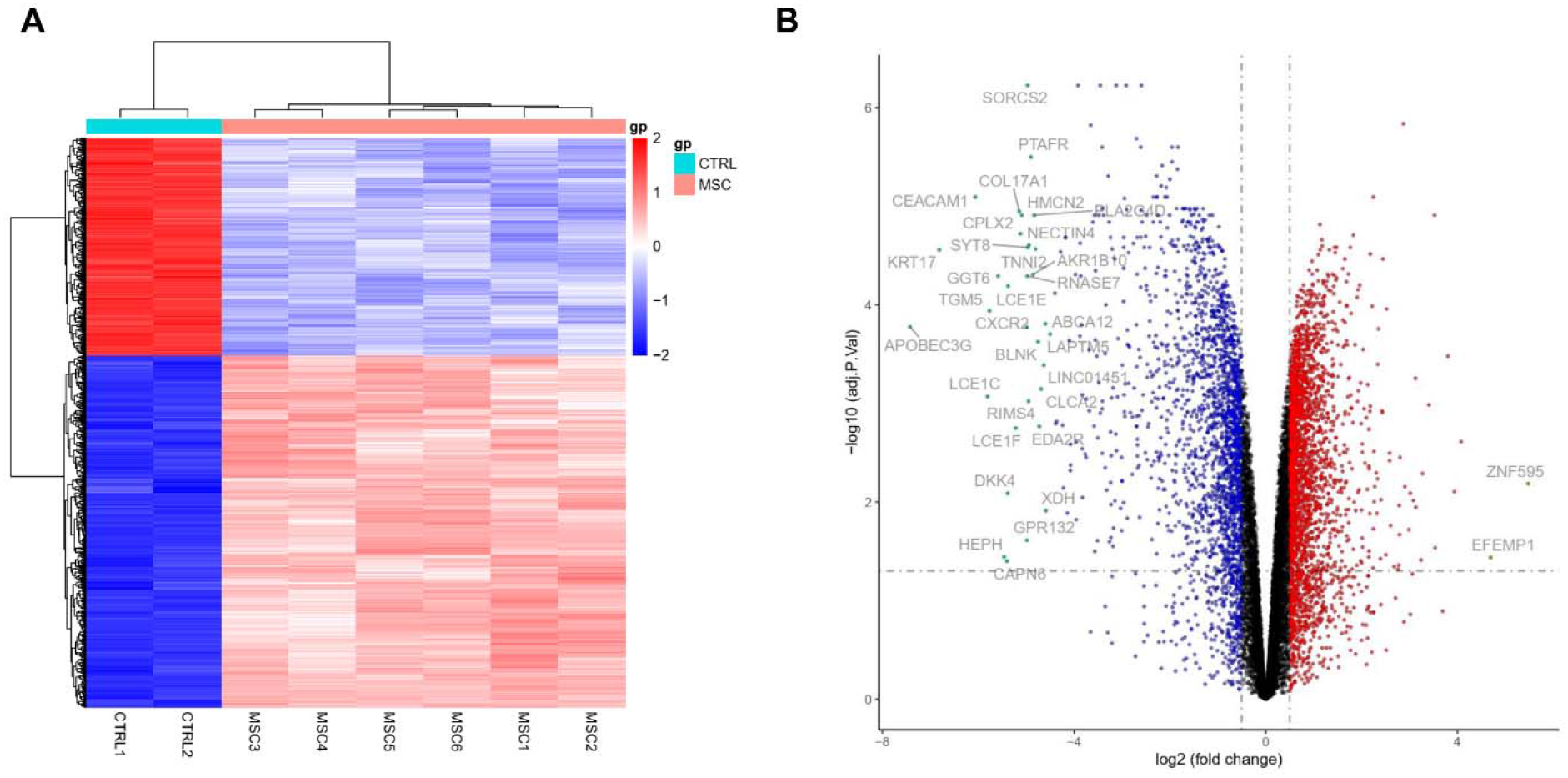
Heatmap and volcano plot of HCT116 cancer cells after DNA damage by irinotecan. (A) Heatmap of significant DEGs. Significant DEGs (P < 0.01) were used to construct the heatmap. (B) Volcano plot for DEGs of HCT116 cancer cells after DNA damage by irinotecan. The most significantly changed genes are highlighted by grey dots.

**Table 1.**

| **Entrez gene** | **Gene symbol** | **Fold-change** | **Regulation** |
| --- | --- | --- | --- |
| Top 10 down-regulated DEGs |  |  |  |
| 60489 | APOBEC3G | -7.424536552 | Down |
| 3872 | KRT17 | -6.811169422 | Down |
| 634 | CEACAM1 | -6.058997335 | Down |
| 9333 | TGM5 | -5.764499265 | Down |
| 124975 | GGT6 | -5.584480624 | Down |
| 353135 | LCE1E | -5.377672374 | Down |
| 1308 | COL17A1 | -5.143544779 | Down |
| 10814 | CPLX2 | -5.1191221 | Down |
| 256158 | HMCN2 | -5.089356824 | Down |
| 3579 | CXCR2 | -4.985685897 | Down |
| Top 10 up-regulated DEGs |  |  |  |
| 7425 | VGF | 3.801006634 | up |
| 79365 | BHLHE41 | 3.523496772 | up |
| 1628 | DBP | 2.874281498 | up |
| 9957 | HS3ST1 | 2.521391179 | up |
| 100533182 | SEC24B-AS1 | 2.443627402 | up |
| 101926975 | LINC01844 | 2.352385673 | up |
| 7857 | SCG2 | 2.338057417 | up |
| 80119 | PIF1 | 2.246041274 | up |
| 100507283 | MSANTD2-AS1 | 2.219586241 | up |
| 6857 | SYT1 | 2.180446776 | up |

### Enrichment analysis of DEGs in colorectal cancer after DNA damage by irinotecan

To further understand the potential mechanisms of DNA damage in cancer cells, we introduced the KEGG and GO analyses (Figure 2). We identified the significant KEGG signaling pathways including “Human papillomavirus infection”, “p53 signaling pathway”, “Axon guidance”, “Hippo signaling pathway”, “Cell cycle”, “Breast cancer”, “Gastric cancer”, “Basal cell carcinoma”, “Melanogenesis”, and “Circadian rhythm”. We also identified the biological processes of GO including “Positive regulation of catabolic process”, “Positive regulation of cellular catabolic process”, “Positive regulation of kinase activity”, “Regulation of autophagy”, “Positive regulation of protein kinase activity”, “Signal transduction by p53 class mediator”, “Gland morphogenesis”, “Positive regulation of autophagy”, “Semaphorin-plexin signaling pathway involved in axon guidance”, and “Semaphorin-plexin signaling pathway involved in neuron projection guidance”. We identified the cellular components of GO “Spindle”, “Organelle outer membrane”, “Outer membrane”, “Mitochondrial outer membrane”, “Site of polarized growth”, “Spindle pole”, “P-body”, “Lysosomal lumen”, “Protein kinase complex”, and “Striated muscle thin filament”. We identified molecular functions of GO including “Catalytic activity, acting on a tRNA”, “Aminoacyl-tRNA ligase activity”, “Ligase activity, forming carbon-oxygen bonds”, “Extracellular matrix binding”, “Frizzled binding”, “Cyclin binding”, “Tropomyosin binding”, “Semaphorin receptor activity”, “Deoxycytidine deaminase activity”, and “Cytidine deaminase activity”.

**Figure 2.**
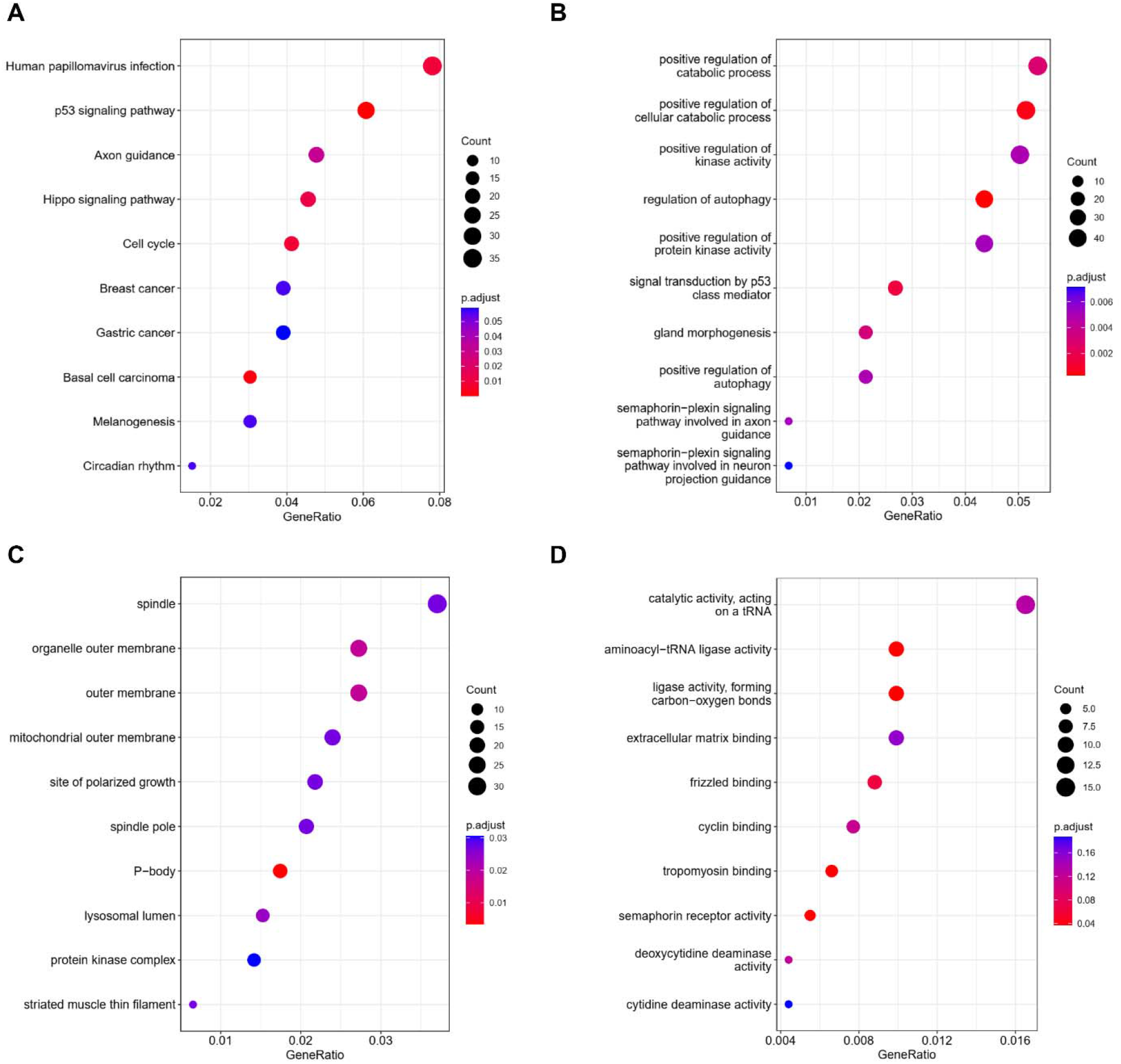
KEGG and GO analyses of DEGs in HCT116 cancer cells after DNA damage by irinotecan. (A) KEGG analysis, (B) Biological processes, (C) Cellular components, (D) Molecular functions.

### PPI network analysis

To explore the relationships among the DEGs, we created the PPI network by using 947 nodes and 3532 edges (combine score > 0.4) with the Cytoscope software. Table 2 indicated the top ten genes with the highest degree scores. The top two significant modules were shown in Figure 3. We further analyzed the PPI and DEGs with Reactome map (Figure 4) and identified the top ten significant processes including “TP53 Regulates Transcription of Cell Cycle Genes”, “TP53 Regulates Transcription of Cell Death Genes”, “Transcriptional Regulation by TP53”, “TP53 Regulates Transcription of Death Receptors and Ligands”, “TP53 Regulates Transcription of Genes Involved in G2 Cell Cycle Arrest”, “TP53 Regulates Transcription of Genes Involved in Cytochrome C Release”, “G0 and Early G1”, “Transcription of E2F targets under negative control by p107 (RBL1) and p130 (RBL2) in complex with HDAC1”, “NR1H2 & NR1H3 regulate gene expression linked to lipogenesis”, and “TP53 Regulates Transcription of Genes Involved in G1 Cell Cycle Arrest” (Supplemental Table S1).

**Figure 3.**
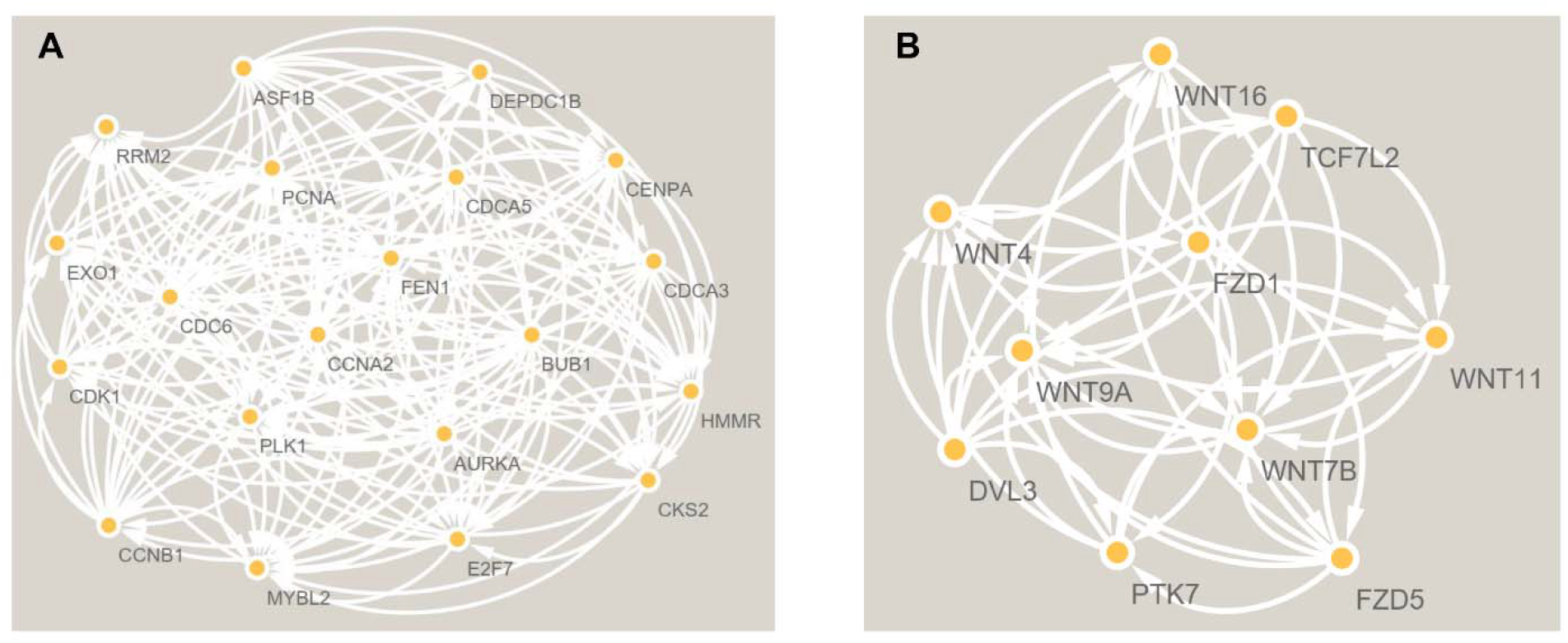
The PPI network analyses of DEGs in HCT116 cancer cells after DNA damage by irinotecan. The cluster (A) and cluster (B) were constructed by MCODE.

**Figure 4.**
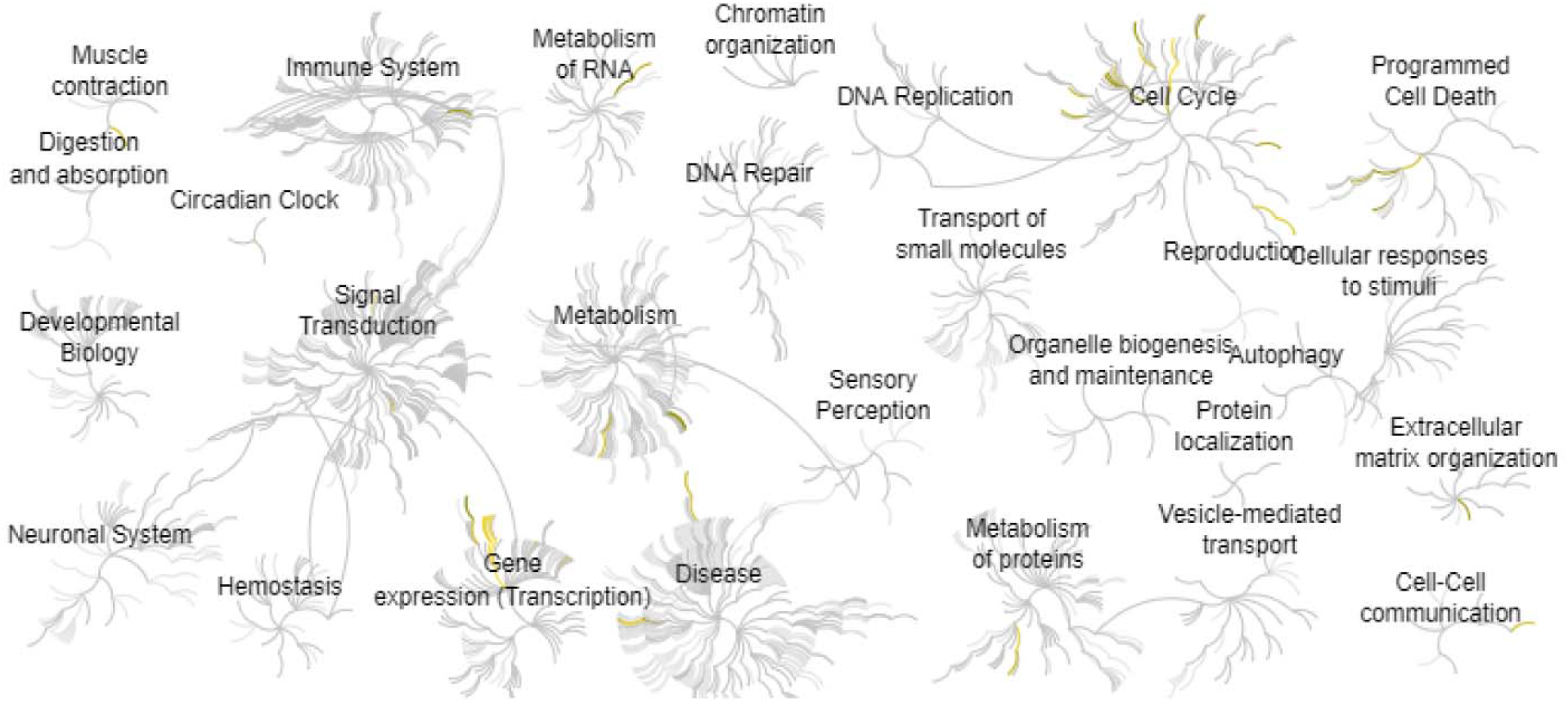
Reactome map representation of the significant biological processes in HCT116 cancer cells after DNA damage by irinotecan

**Table 2:**
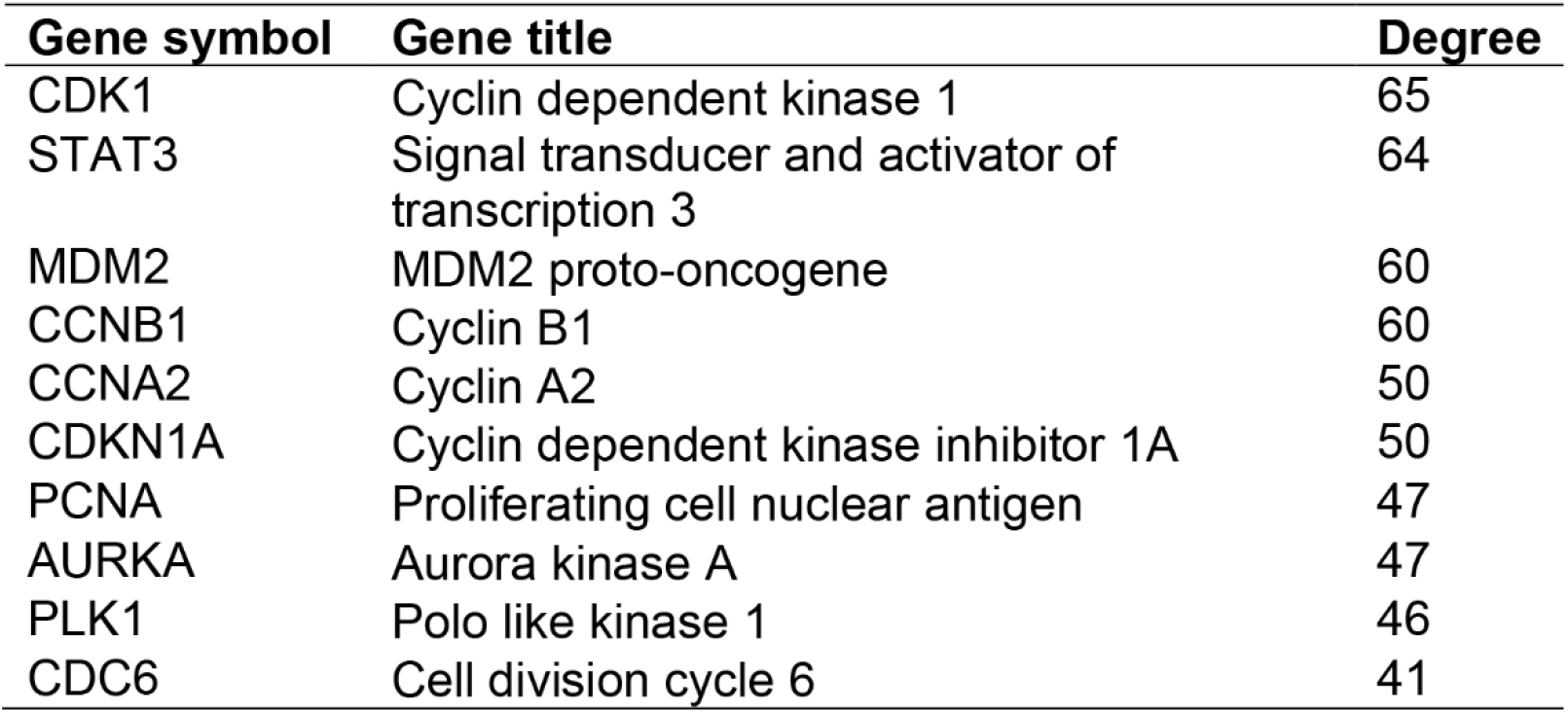
Top ten genes demonstrated by connectivity degree in the PPI network.

## Discussion

Colorectal cancer is the third most common cancer in the US and the third most reason for cancer-related death^15^. Targeted therapy is to use the biological cancer features such as immune cells and other cells to inhibit the progression of cancer^16^. However, it is difficult to completely kill the cancer cells due to the complex microenvironment. This study is to help understand the complex biology of colorectal cancer through genomic analysis.

Based on our analysis, we found that the p53 signaling pathway and Hippo signaling pathway are major affected processes in colorectal cancer by DNA damage. As a tumor suppressor, p53 is majorly found in colorectal cancers. Hak-Su Goh et al found cancer patients with p53 mutation showed a poorer prognosis than those patients without mutations. Moreover, cancer patients showed responses to therapies are largely dependent on p53 mutations^17^. Shuxia Liu et al also found the anti-p53 antibody in the blood can be considered a diagnostic biomarker for colorectal cancer, which has the potential to distinguish the context of colorectal cancer from healthy controls or benign diseases^18^. The hippo pathway contains a number of proteins that regulate the growth of different tissues during development and regeneration such as cancer. The deregulation of the Hippo pathway has been shown in a wide range of different cancers including lung, colorectal, liver and prostate cancers, and it is closely related to the poor prognosis^19^. Besides these significant signaling pathways, the circadian rhythm is also indicated a great impact during the progression of colorectal cancer. Circadian clocks regulated various physiology to maintain homeostasis such as metabolism, immune, and senescence^20–30^. Disruption of the circadian clocks plays essential roles in downstream targets and diseases including cancers. Recently finding showed targeting the core clock genesis a novel approach in cancer therapy^31^.

In this study, we also discovered the interactive molecules during DNA damage-induced colorectal cancer. J K Buolamwini et al found that CDK1 inhibitors can control the cell cycle in cancers, which has been demonstrated in preclinical studies and clinical trials^32^. Yan Lin et al found the STAT3 and IL6 signaling in colorectal cancer show an effect on tumor-infiltrating immune cells in the tumor microenvironment^33^. GPCR and its related-RGS signaling pathways are associated with several critical human diseases including cancer, arthritis, and aging-related diseases^34-43^. Stéphane Pelletier et al found GPCR agonists stimulate tyrosine phosphorylation of STAT3 proteins in a Rac-dependent manner, which is critical in the development of cancers^36^. Rami M Elshazli et al found MDM2 (rs2279744) indicated relationships with colorectal cancer risks among Asians and Africans under a recessive model by stratification analysis^44^. Gennadi V Glinsky et al found cancer cells indicate a stem cell-like expression profile such as increased levels of certain cell cycle marker proteins CCNB1 and CCNA2^45^. Kaori Shima et al found CDKN2A promoter methylation and gene silencing are related to the CpG island methylator phenotype in colorectal cancer^46^. Recently, targeting PCNA is considered to be as an effective method to repress the proliferation of cancer cells^47^. AURKA is a serine/threonine kinase, which shows higher expression in cancer tissues according to TCGA database^48^. Xiang Ding et al found PLK1 is a critical gene in colorectal cancer, which is positively correlated with the development of cancer^49^. Chang Yu et al found CDC6 may serve as a prognostic biomarker in colorectal cancer^50^.

In summary, our study discovered the significant effects of DNA damages on colorectal cancer. The P53 signaling pathway and Hippo signaling pathways are the major affected processes during the development of colorectal cancer. Our study provides valuable insights for the treatment of colorectal cancer.

## Author Contributions

Hongmei Guo, Mason Zhang: Methodology and Writing. Hanming Gu, James Liu: Conceptualization, Writing-Reviewing and Editing.

